# Root Contours Guided Design of a Multicellular PID Controller

**DOI:** 10.1101/2025.05.05.652201

**Authors:** Vittoria Martinelli, Davide Fiore, Davide Salzano, Mario di Bernardo

## Abstract

Ensuring a stable and robust phenotype expression is a key challenge for the correct operation of synthetically engineered cells, and feedback has been highlighted as a key mechanism to achieve this goal. Biomolecular PID controllers have been extensively leveraged at a single cell level to regulate gene expression. However, single-cell architectures suffer from limited modularity and might pose a significant challenge for their in vivo implementation due to high metabolic load and possible incompatible reactions. To overcome these limitations, it has been proposed to distribute the control actions over different cell populations realizing a multicellular feedback control architecture. In this paper we provide design guidelines derived by means of the root contours method to tune the control gains of a multicellular PID controller. We then validate performance, robustness and modularity of the multicellular PID controller through in silico simulations in BSim.

## I. INTRODUCTION

Synthetic biology integrates engineering principles into the design of genetic circuits to endow cells with new functionalities. Applications of synthetically engineered cells range from microbial diagnostics and medical treatments [1], to biofuels made with engineered microorganisms [2], and innovative food production tailored to meet specific nutritional needs [3]. However, the inherently nonlinear and stochastic nature of the biochemical processes involved poses significant challenges for the design of robust circuits able to maintain the desired phenotype despite disturbances.

*Cybergenetics* addresses these challenges designing synthetic feedback control architectures within cells [4]. Although the control literature is ripe with solutions providing high closed-loop performance, constructing these advanced strategies within cells is not possible due to the limited availability of suitable biological components. Therefore, the implementation of simple yet reliable controllers such as Proportional-Integral-Derivative (PID) controllers has been preferred [5], [6]. Biomolecular PID controllers can indeed ensure precise regulation [7] and provide a fast and damped response through the derivative action. The biomolecular PID proposed in the literature requires embedding all the control functionalities within the same cell, which poses significant implementation challenges due to factors such as excessive metabolic burden and lack of modularity. To tackle these problems, a promising solution is to distribute the control components across different cell populations within a microbial consortium [8], alleviating the burden on each cellular population and increasing the modularity. In this architecture, the control actions realizing the PID control are distributed in different cell populations and can be combined to realize P, PI [9], PD [10], and PID [11] controllers.

In this paper, after reviewing the multicellular PID control architecture presented in [11], we derive analytical conditions on the biological parameters of the control architecture that can be used to choose the control gains so as to fulfill a set of given design requirements on the static and transient performance. The conditions are derived by means of the root contours method, which allows us to obtain relationships between the control gains and the performance of the closed-loop system locally to the desired equilibrium point. We compare the performance and robustness of the multicellular P, PI, PD and PID control architectures through *in silico* experiments carried out in BSim, an agent-based environment designed to simulate bacterial populations [12].

## II. MULTICELLULAR PID CONTROL ARCHITECTURE

The multicellular Proportional-Integral-Derivative (PID) control consists in three controller cell populations implementing the biological equivalent of the classical PID control law

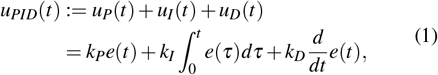

where *k*_*P*_, *k*_*I*_ and *k*_*D*_ are the proportional, integral and derivative control gains, respectively, and *e*(*t*) is the control error, defined as *e*(*t*) := *y*_d_(*t*) − *y*(*t*), with *y*_d_(*t*) the desired output, and *y*(*t*) being the measured output of the controlled process Φ(*t*) embedded in a fourth population denoted as targets (Fig. 1.a). The overall control signal *u*_*PID*_(*t*) computed by the controllers is sensed by the target population hosting the process Φ(*t*), whose output *y*(*t*) is fed back to the controllers closing the control loop. The communication between controllers and targets is realized by the pair of orthogonal quorum sensing molecules, *Q*_*u*_ and *Q*_*x*_, that act as proxies of the control input *u*_*PID*_(*t*) and of the process state *y*(*t*), respectively. These molecules are produced by the cells and diffuse through their membranes into the environment. In what follows we use the superscripts *t, p, i, d* and *e* to denote quantities in the targets, in the proportional, integral and derivative controller cells, and in the external environment, respectively.

**Fig. 1:**
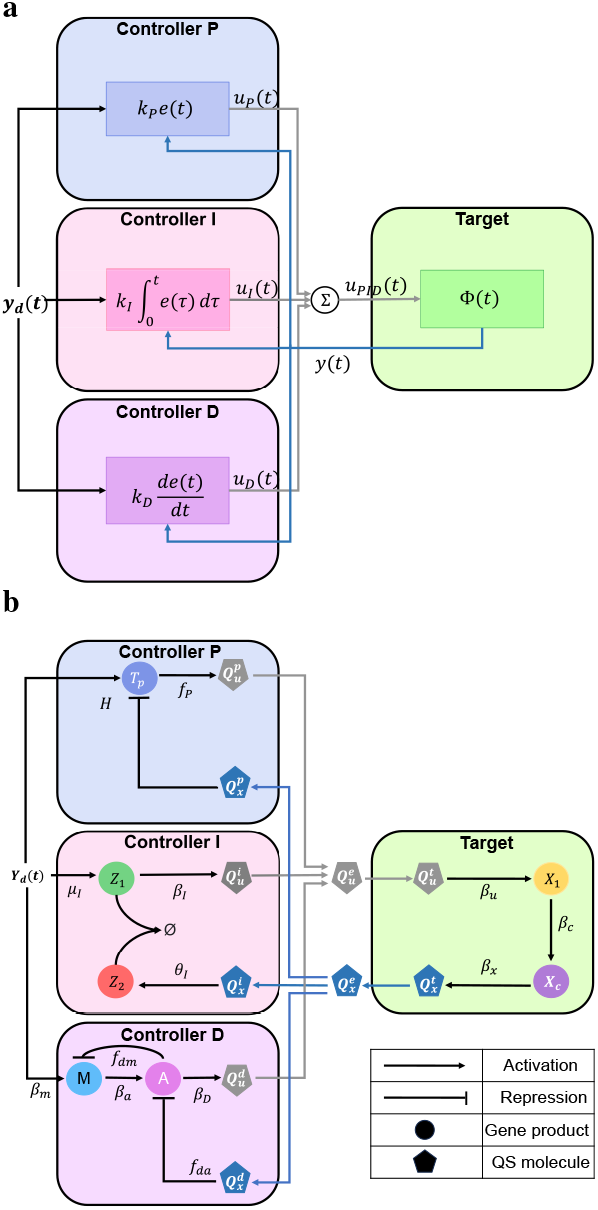
Multicellular PID control architecture. (a) The proposed control strategy involves three controller populations comparing the reference *y*_*d*_ (*t*) to the output *y*(*t*) of the controlled process Φ(*t*) embedded in the target population. Based on this mismatch, they collectively compute the overall control signal *u*_*PID*_(*t*), closing the feedback loop. (b) The four cell populations exchanging biomolecular signals via two orthogonal quorum sensing molecules. Specifically, the controller cells produce the control molecule *Q*_*u*_ based on the mismatch between the reference signal *Y*_d_ and the measure of the output *X*_*c*_ of the controlled process. This measure is broadcast to the entire consortium via the sensor molecule *Q*_*x*_, produced by the target cells to close the feedback loop.

### A. Aggregate dynamics

The aggregate model describes the evolution of the concentration of each biochemical species averaged over the whole microbial consortium. Details on the derivation of the aggregate dynamics from the agent-based dynamics can be found in [11]. The model is derived under three simplifying assumptions: i) all cells grow and divide at the same rate, implying that each chemical species is diluted at the same rate *γ*; ii) the consortium is balanced, meaning that all populations have the same number of cells *N*; iii) the quorum sensing molecules diffuse with a uniform diffusion rate *η*.

The dynamics of the species *X*_1_ and *X*_*c*_ in the targets (Fig. 1.b, green cell) [5], [7], can be modeled as

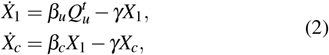

where *β*_*u*_ and *β*_*x*_ are the activation rates of *X*_1_ and *X*_*c*_, respectively. Note that the production of *X*_1_ is promoted by 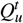, which is produced by the controller populations. Additionally, the targets produce an orthogonal quorum sensing molecule *Q*_*x*_, whose production is proportional to *X*_*c*_. This is used as a proxy for the concentration of *X*_*c*_. The dynamics of *Q*_*x*_ within the target cells can be formally described by

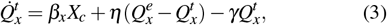

where *β*_*x*_ is the activation rate of *Q*_*x*_.

Each controller population contributes to the production of the control molecule *Q*_*u*_. Specifically, the proportional controller cells (light blue in Fig. 1) embed a network inspired by the design presented in [5], where two transcription factors compete for the promoter producing the quorum sensing molecule 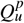. Its dynamics is given by [11]

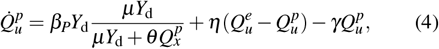

where *β*_*P*_ plays the role of a proportional gain, and *μ* and *θ* are control parameters. The first term in (4) can be proven to be a nonlinear function of the control error, defined as *e*(*t*) := *μY*_d_ −*θ Q*_*x*_, where *y*_d_ = *μY*_d_ is the desired output and *y*(*t*) = *θ Q*_*x*_(*t*) is the measured output [5].

The integral controller population (light red in Fig. 1) contains a network based on the antithetic motif introduced in [7], comprising two proteins, *Z*_1_ and *Z*_2_, that can bind forming an inert complex. The dynamics of such species is described by

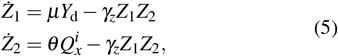

where *μ* and *θ* are the activation rates of *Z*_1_ and *Z*_2_, respectively, and *γ*_*Z*_ is their annihilation rate. The difference (*Z*_1_ − *Z*_2_) is proportional to the integral of the control error *e*(*t*), thus implementing an integral control [7]. The integral action is delivered via the control molecule 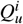, which is produced proportionally to *Z*_1_, following

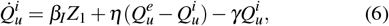

where *β*_*I*_ acts as an integral control gain. The last contribution to the control action is provided by the derivative controller population (purple in Fig. 1). These cells contain a circuit based on the synthetic network presented in [5], whose dynamics can be written as

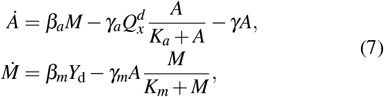

where *β*_*a*_ and *β*_*m*_ are the activation rates of *A* and *M*, respectively. In addition, the active degradation of the two biomolecular species is modeled through an enzymatic reaction based on the Michaelis-Menten function characterized by constants *K*_*a*_ and *K*_*m*_, and rates *γ*_*a*_ and *γ*_*m*_. Under the assumption of enzyme saturation, i.e. *K*_*a*_ ≪ 1 and *K*_*m*_ ≪ 1, it can be shown that 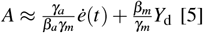. In these cells the control molecule 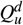 is produced proportionally to the concentration of *A*. Formally, its dynamics is described by

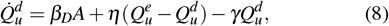

where *β*_*D*_ plays the role of a derivative control gain.

Finally, the dynamics of the molecules diffusing into the cells where they are not produced can be described as

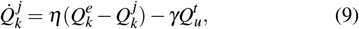

where *k* ∈ {*x, u*} and *j t*, ∈ {*p, i, d*}, and the dynamics of *Q*_*u*_ and *Q*_*x*_ in the external environment can be written as

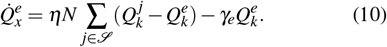

Here, *N* is the number of cells in each population, and ℐ = {*t, p, i, d*}. Note that, the mathematical models of the multicellular P, PI and PD controllers derived in [9], [10] can be directly obtained from this model.

### B. Simplified dynamics

To obtain meaningful conditions that can be used to tune the control gains, the above model can be further simplified under the following assumptions [11]:

**A1**: *The quorum sensing molecules diffuse at a rate that is considerably faster than their degradation, i*.*e. η* ≫Γ_*PID*_, *where* Γ_*PID*_ := 4*γ. This assumption has been validated in various models parameterized based on in vivo experiments [13]*.

**A2**: *The sequestration rate between species Z*_1_ *and Z*_2_ *is sufficiently rapid, i*.*e. Z*_1_ ≫ *Z*_2_ *at steady state*.

**A3**: *The enzymatic reactions described in* (7) *occur under substrate saturation, i*.*e. K*_*a*_ ≫ *A and K*_*m*_ ≫ *M [5]*.

**A4**: *The dynamics of the derivative network in* (7) *are significantly faster than those of the target process in* (2), *enabling a time-scale separation between the controlled process dynamics and the derivative cells dynamics (details can be found in [5]*).

If assumptions **A1**-**A4** hold, the dynamics of the closed loop architecture can be reduced to

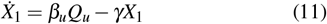

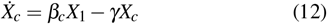

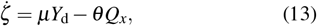

where *ζ* := *Z*_1_ − *Z*_2_, and 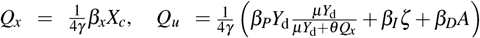 and 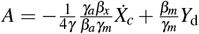 are the steady-state values of the sensor molecule *Q*_*x*_, the control molecule *Q*_*u*_ and the species *A*, respectively. See [11] for more details.

## III. ARCHITECTURE DESIGN

In this section, we will derive analytical conditions on the biological parameters of the multicellular PID architecture that can be used to tune the control gains. Specifically, the conditions will be derived by means of the root contour method [14], which allows to study how the value of the control gains affect the position in the complex plane of the poles of the transfer function of the closed loop systems linearized around the desired set-point. Note that, the underlying assumption is that there exist a unique, globally asymptotically stable equilibrium point in the positive orthant. This analysis can be employed to tune the control gains so that, locally to the equilibrium point, the output of the target cells is regulated to the desired set-point with prescribed static and transient performance, defined in terms of the steady-state error *e*_∞_ := lim_*t*→∞_ |*e*(*t*)|, settling time 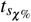, and overshoot *o*_%_.

### A. Root contours guided design

We derived the set-point from the reduced model (11)-(13), that is, 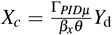. This equilibrium can be shown to be admissible only if [11]:

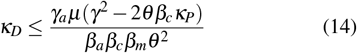

Next, we linearized the system around this set-point and derived, in the Laplace domain, the closed-loop transfer function between the input *μY*_d_(*s*) and the output *θ Q*_*x*_(*s*).

This function can be computed to be

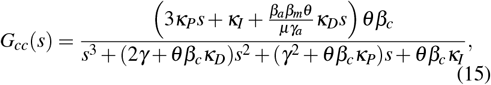

where the control gains 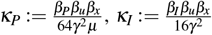 and 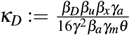 come from the aggregation of system parameters (see [11] for details on the derivation).

We first consider the static gain of *G*_*cc*_(*s*) in Eq. (15), which represents the ratio between the output and the input under steady-state condition. This gain, computed as *s* → 0 (*t* → ∞), is always equal to 1 if *κ*_*I*_ ≠ 0. This means that output matches the input, resulting in a steady-state error equal to zero. However, this does not always hold when *κ*_*I*_ = 0. For example, for the proportional controller alone (i.e. *κ*_*D*_ = *κ*_*I*_ = 0), 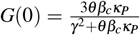 equals 1 only if *κ*_*P*_ is precisely tuned to a specific value based on the nominal parameters. Thus, robust asymptotic regulation can be guaranteed only when the integral action is employed.

To obtain analytical relationships between the control gains and the closed-loop performance, we applied the root contours on Eq. (15). This graphical technique generalizes the root locus method by investigating how multiple parameters influence the closed-loop response [14]. Specifically, the root locus of a loop transfer function *L*(*s*) is the set of points where 1 + *ρL*(*s*) = 0, with *ρ* being a control parameter.

To apply the root contours on *G*_*cc*_(*s*) in Eq. (15), we first set *κ*_*D*_ = *κ*_*I*_ = 0 and then recast the denominator of Equation (15) as:

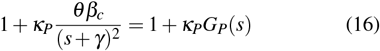

where 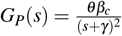 is the loop transfer function of targets controlled by only a Proportional controller population, as shown in [9]. The points belonging to the root locus of *G*_*P*_(*s*) are shown as the dashed light gray vertical line in Fig. 2.

**Fig. 2:**
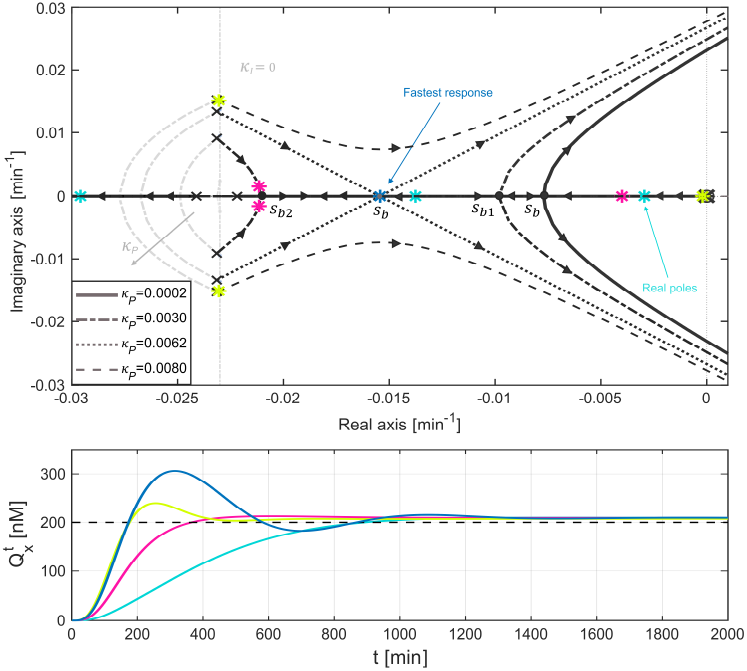
Tuning of the control gains using the root contours method. The upper panel shows the root contours, while the lower panel shows the time evolution of the concentration of the measured output 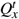 for fixed control gain values. In the upper panel, the dashed light gray lines represent the root contours of the P and PD controllers (for *κ*_*I*_ = 0), while the black arrows indicate increasing values of *κ*_*I*_. After selecting admissible values of *κ*_*P*_ and *κ*_*D*_ (see condition (14)), we set some values of *κ*_*I*_ fulfilling (20). Specifically, the three coincident real poles (blue star) are placed by choosing *κ*_*I*_ meeting also condition (24), ensuring the fastest possible response (blue line, lower panel). The other biochemical parameters were selected as in Appendix A, whereas the reference signal was set to *Y*_d_ = 60 nM.

Next, fixing 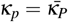 and setting *κ*_*D*_ ≠ 0, *κ*_*I*_ = 0, we recast the denominator of Eq. (15) as:

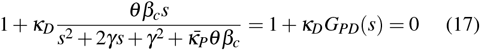

where 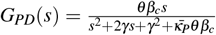 is the loop transfer of the multicellular PD controller described in [10]. Here, for any fixed value of *κ*_*P*_ it is possible to draw the root locus of *G*_*PD*_(*s*). The starting poles of the locus (i.e. the position of the poles when *κ*_*D*_ = 0) are selected by choosing the points of the root locus of *G*_*P*_(*s*) corresponding to 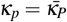. Different root loci associated to different values of *κ*_*P*_ are illustrated in Fig. 2 as dashed light gray lines.

Finally, the root contours of the full PID architecture was derived by setting 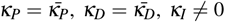 and recasting the denominator of Equation (15) as

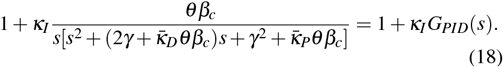

Here, 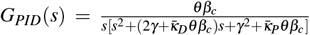 represents the loop transfer function of the PID control architecture and *κ*_*I*_ is selected as the variable parameter. The root contour of *G*_*PID*_(*s*) has as starting poles (i.e. the position of the poles when *κ*_*I*_ = 0) the points lying on the root contour of *G*_*PD*_(*s*) when 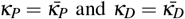. Specifically, each root contour has three branches originating at:

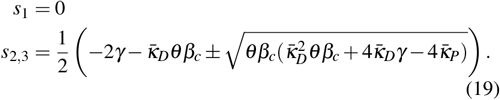

These branches approach infinity as *κ*_*I*_ →∞ (see black lines in the upper panel of Fig. 2). As such, increasing *κ*_*I*_ can destabilize the closed loop system. To ensure stability it is necessary that

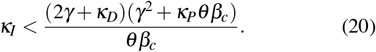

Eq. (20) shows that choosing fast-diving cells or reducing the strength of the promoter induced by *Q*_*x*_ in the integral cells can expand the range of the possible values for *κ*_*I*_.

Furthermore, condition (14) constrains the initial points from which the branches of the root contours of *G*_*PID*_(*s*) originate. Specifically, the starting points *s*_2,3_ can either be both real or complex conjugates. All the other conditions on the control parameters are discussed in greater detail in [11].

### Real starting points

When the integral gain *κ*_*I*_ = 0, the poles *s*_2,3_ from Eq. (19) are real only if the following condition holds (see Fig. 2 for *κ*_*P*_ = 0.0002, light blue stars):

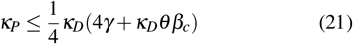

As *κ*_*I*_ increases, the closed-loop poles remain real until the breakaway point *s*_*b*_, where two real branches become complex. This point is characterized by the most negative real part of the dominant pole of the closed loop system, which corresponds to the fastest response. As such, given that 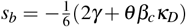, by selecting 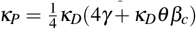 and choosing the integral gain as

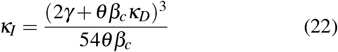

it is possible to obtain the best performance in terms of both percentage overshoot and settling time.

### Complex conjugates starting points

The poles *s*_2,3_ are complex conjugates if (Fig. 2 for *κ*_*P*_ = 0.0030, pink stars):

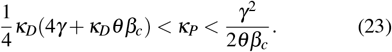

Increasing the value of *κ*_*I*_, the complex branches (and thus the closed-loop poles) become real in the breakaway point *s*_*b*1_, and then become complex again after the breakaway point *s*_*b*2_. These two breakpoints overlap in *s*_*b*_ if 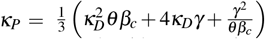, as shown in Fig. 2 for *κ*_P_ = 0.0062, blue coincident stars. The leftmost value of *s*_*b*_, which corresponds to the fastest closed-loop system, is 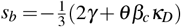 for *κ*_*I*_ selected as

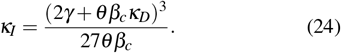

Finally, for 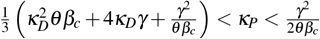 there are always two complex conjugate poles (see [11]). In this scenario, the best overshoot and settling times can be obtained when the real part of the complex poles is closest to zero, which happens when condition (24) holds. Note that, despite being local to the equilibrium point, our analysis is relevant to synthetic biology applications as these circuits are typically designed to function around the set point [15].

A similar analysis can be conducted to study the local transient performance of the P, PI and PD control strategies presented in [9], [10]. In Fig. 2 (bottom panel) we show numerical simulations in Matlab on the nonlinear system, where we selected the control gains so as to achieve the lowest overshoot and the fastest settling time, while ensuring error equal to zero at steady state.

## IV. IN SILICO EXPERIMENTS

We present *in silico* experiments to confirm the effectiveness of the proposed multicellular PID architecture, and compare its performance with those of the other architectures within the PID family in more realistic conditions. First, we test the sensitivity to possible uncertainty in the estimation of target cell’s parameters, which can be due to the high variability naturally exhibited by biological systems. Next, we assess sensitivity to cell-to-cell variability, an intrinsic property of natural biological systems. Finally, we test robustness to population imbalances due to unavoidable asymmetries in the metabolic load on each population.

We performed the *in silico* experiments in Matlab or BSim, an agent-based realistic simulator designed for bacterial populations [12]. This simulation platform is designed to account for cell growth and division, diffusion of molecules and cell-cell physical interactions; additionally, it allows to change the geometry of the hosting chamber. Specifically, we set up a scaled-down version of the BSim growth chamber used in [16], with dimensions 23 *μ*m × 15 *μ*m × 1 *μ*m, hosting around 120 cells, to reduce computational time while maintaining statistical significance.

We specified the control objective as that of achieving a steady-state error *e*_∞_ ≤ 0.1, with a settling time less than 24 hours, that is, 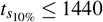 min. Furthermore, we required any oscillations to vanish at steady state, with an overshoot *o*_%_ ≤ 20%. To highlight modularity, we used the same control gains across all control architectures, selected using the root contours method so as to meet the desired control criteria.

First, we ran numerical experiments in Matlab where targets had parameters differing from the one used in the design step. This mimics uncertainty in the estimation of cell’s parameters. In these simulations we neglected cell growth and division and the spatial effects due to molecules diffusion, in order to minimize the computational effort. Figure 3 shows that on average all control architectures can reduce the error to zero, regulating *Q*_*x*_ to the desired value. However, only by including an integral action ensures that the steady state error is independent on the choice of *κ*_*P*_ and *κ*_*D*_ (provided that (14) is satisfied). In contrast, employing an integral action results in a worse settling time. More precisely, we observed an average settling time of 438 min for the PI architecture and 478 min for the PID architecture, and of 327 min and 183 min for P and PD strategies, respectively.

**Fig. 3:**
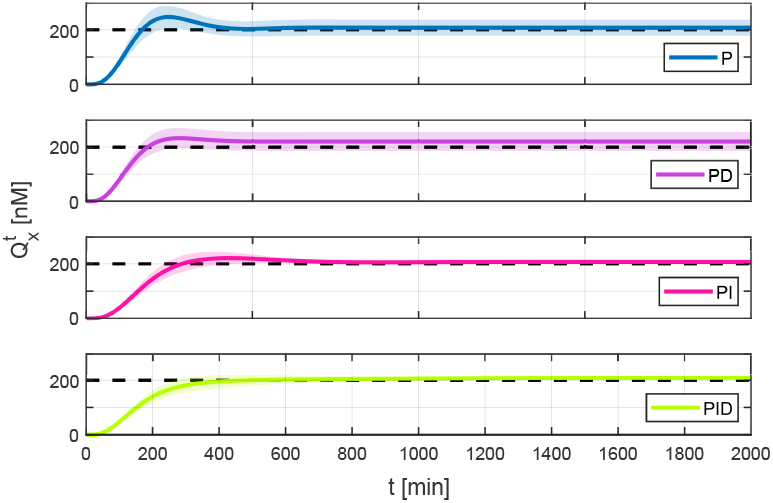
*In silico* regulation experiments in Matlab. Mean (solid line) and standard deviation (shaded area) of 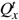 when the targets are controlled by P, PD, PI and PID control actions, respectively. Control gains were tuned to meet condition (14) and from the regions shown in Fig. 2, fulfilling the control requirements on the transient response, and their values kept the same for all strategies to exploit modularity. In each of the 10 simulations, the targets’ parameters were drawn from a normal distribution centered at their nominal value 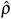 with standard deviation 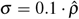 to test the robustness to uncertainties in the biological process Φ(*t*). All the other parameters were fixed to their nominal values as in Appendix A.

Next, we tested the robustness of the control architectures in BSim. Specifically, we assessed the sensitivity to cell-to-cell variability, that is, differences in expression levels among individual cells within a population caused by eventual uneven distribution of the genetic material between cells after division. We evaluated the performance in terms of settling time, overshoot and percentage steady-state error, defined as 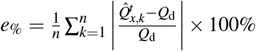. Here, 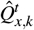 is the value of 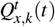, averaged over the last 600 min of the *k*-th experiment, 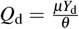 is the desired value of *Q*_*x*_, and *n* is the total number of experiments at a given heterogeneity level to simulate cell-to-cell variability. We assigned different values of the targets’ and controllers’ parameters at each cell division, drawing them from a normal distribution with mean their nominal value 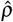 and standard deviation 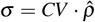, where *CV* stands for the coefficient of variation. Fig. 4a shows that all the control strategies fulfill the desired control requirements, except for the P controller when *CV* = 0.15, where it exceeds the upper bound of 20%. Additionally, the metrics of each controller remains almost the same for all the values of *CV*, except for the PI and PID when *CV* = 0.15, where the settling time increases of about 2.5 times and 1.2 times, respectively, with respect to when *CV* = 0.

**Fig. 4:**
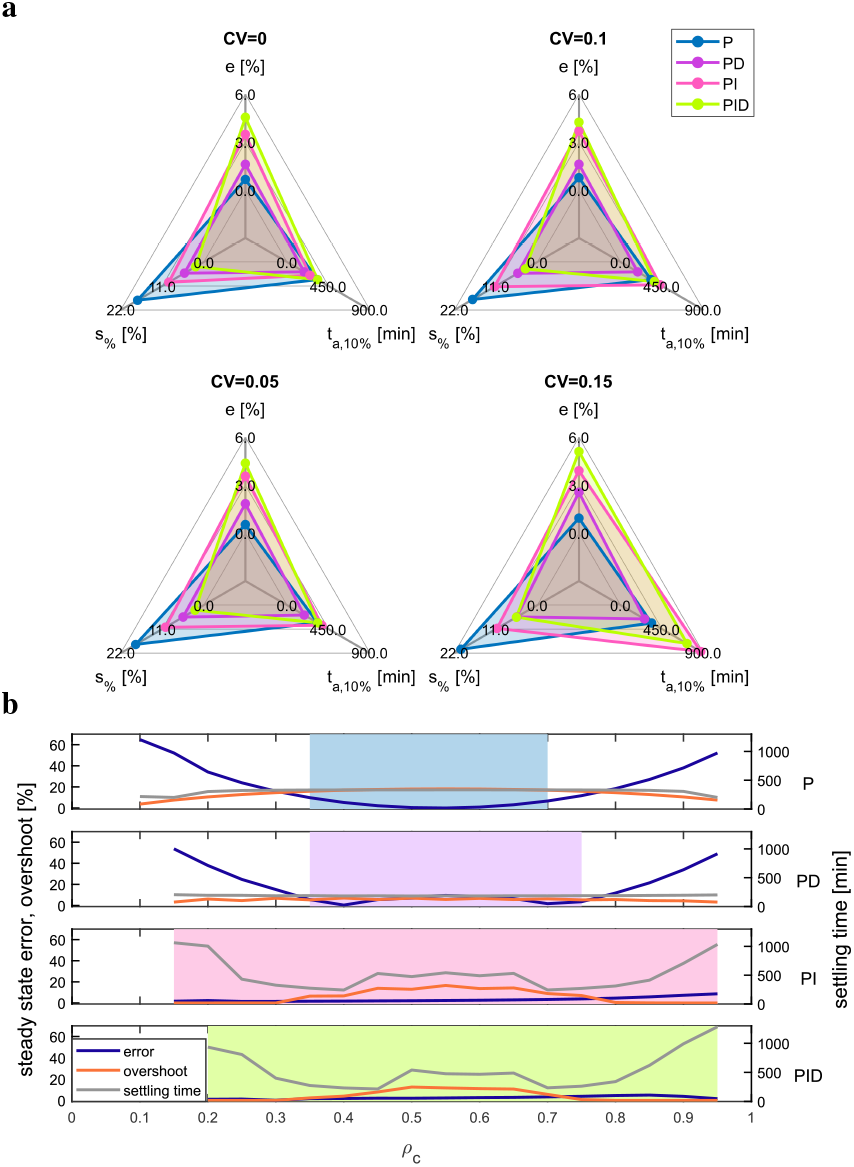
Robustness of the P, PD, PI and PID control architectures. (a) Percentage error at steady state, settling time and overshoot as parameter variations increase. For each value of the *CV* ∈ {0.05, 0.1, 0.15} we run *n* = 20 experiments, drawing the controllers’ and targets’ parameters from a normal distribution centered at their nominal value 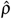 and with standard Deviation 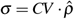. The BSim microfluidic chamber varied across control strategies to maintain the same targets’ number at steady-state. The metrics were computed filtering the average targets’ output signal with a moving average filter of window width *W* = 60. The initial number of cells for each control strategy was equally divided between the different cell types. (b) Steady-state percentage error, settling time and overshoot ratio of the controller cells *ρ*_*c*_ := *N*_*c*_*/N* varies, with *N* = 20 cells in the chamber. The growth rate was set to zero. The shaded areas evidence the values of *ρ*_*c*_ corresponding to successful simulations meeting control requirements. In all experiments the reference signal was *Y*_d_ = 60 nM, while the gains were chosen as *β*_*P*_ = 0.0414 min^−1^, *β*_*D*_ = 0.0933 min^−1^ and *β*_*I*_ = 0.0002 min^−1^. The value of the parameters can be found in Appendix A.

Finally, we assessed the robustness to consortium composition imbalances, which can stem from metabolic load asymmetries. To this aim, we neglected cell growth and division and evaluated the control metrics mentioned above when different members of controllers and targets are present in the consortium. Here, we defined the residual steady state error as 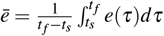, where *t*_*s*_ = 1440 min, and *t*_*f*_ is the simulation time set to be 2000 min. For each experiment we varied the percentage of controller cells *ρ*_*c*_ := *N*_*c*_*/N*, with *N* the total number of cells in the chamber, assuming all controller populations are in the same number.

We observed that the integral action ensure robustness even for high population imbalances, ensuring that both static and dynamic requirements are met even in the presence of extreme imbalances (see shaded areas in Figure 4). Instead, Proportional and PD controllers can meet the specifications only when the imbalance between targets and controllers is limited.

## V. Discussion

In this paper we presented a method to derive analytical conditions for tuning the control gains of a multicellular PID control architecture to meet desired performance. Building on the multicellular PID framework introduced in [11] and after deriving the aggregate dynamics of the consortium, we provided some guidelines to tune the control gains using the root contours method. Numerical experiments confirmed the effectiveness of the control strategy, showing that the integral action ensures robust regulation, while adding derivative action enhances the transient response. The approach presented here is not limited to the multicellular consortium illustrated above, but to any multicellular architecture whose closed-loop stability and performance depend on two or more tunable parameters. A key open problem for the *in vivo* deployment of this architecture is to devise strategies that ensure long-term coexistence of cell populations in the consortium, potentially through embedded circuits, external controllers, or multi-chamber bioreactors [17], [18].

## Appendix

### A. Nominal biochemical parameters

The growth and mechanical parameters used in the BSim simulations were selected as in [13], while the nominal biochemical parameters were chosen as: *β*_*u*_ = 0.06 min^−1^, *β*_*x*_ = 0.03 min^−1^, *γ* = 0.023 min^−1^, *η* = 2 min^−1^ (from [13]); *β*_*c*_ = 0.1 min^−1^, *μ* = 1 min^−1^, *θ* = 0.3 min^−1^, *γ*_*z*_ = 0.01 nM^−1^min^−1^, *β*_*a*_ = 1.5 min^−1^, *β*_*m*_ = 0.4167 min^−1^, *γ*_*a*_ = *γ*_*m*_ = 1.5 min^−1^ (from [5]); *γ*_*e*_ = 0.0023 min^−1^.

## References

[1] Z. Zhou, et al., “Engineering probiotics as living diagnostics and therapeutics for improving human health,” Microbial Cell Factories, vol. 19, no. 1, 2020.

[2] Z. Liu, et al., “Yeast synthetic biology advances biofuel production,” Current Opinion in Microbiology, vol. 65, pp. 33–39, 2022.

[3] S. Shi, et al., “Synthetic biology: a new frontier in food production,” Trends in Biotechnology, vol. 40, no. 7, pp. 781–803, 2022.

[4] M. H. Khammash, “Cybergenetics: Theory and applications of genetic control systems,” Proceedings of the IEEE, vol. 110, no. 5, pp. 631– 658, 2022.

[5] M. Chevalier, et al., “Design and analysis of a proportional-integral-derivative controller with biological molecules,” Cell Systems, vol. 9, no. 4, pp. 338–353.e10, 2019.

[6] M. Filo, et al., “Biomolecular feedback controllers: from theory to applications,” Current Opinion in Biotechnology, vol. 79, p. 102882, 2023.

[7] C. Briat, et al., “Antithetic integral feedback ensures robust perfect adaptation in noisy biomolecular networks,” Cell Systems, vol. 2, no. 1, pp. 15–26, 2016.

[8] I. Ruolo, et al., “Control engineering meets synthetic biology: Foundations and applications,” Current Opinion in Systems Biology, vol. 28, p. 100397, 2021.

[9] V. Martinelli, et al., “Multicellular PI control for gene regulation in microbial consortia,” IEEE Control Systems Letters, vol. 6, pp. 3373– 3378, 2022.

[10] V. Martinelli et al., “Multicellular PD control in microbial consortia,” IEEE Control Systems Letters, vol. 7, pp. 2641–2646, 2023.

[11] V. Martinelli, et al., “Multicellular PID control for robust regulation of biological processes,” Journal of The Royal Society Interface, vol. 22, 2025.

[12] A. Matyjaszkiewicz, et al., “Bsim 2.0: An advanced agent-based cell simulator,” ACS Synthetic Biology, vol. 6, no. 10, pp. 1969–1972, 2017.

[13] D. Fiore, et al., “Multicellular feedback control of a genetic toggleswitch in microbial consortia,” IEEE Control Systems Letters, vol. 5, no. 1, pp. 151–156, 2021.

[14] F. Golnaraghi et al., Automatic control systems. McGraw-Hill Education, 2017.

[15] N. Olsman, et al., “Architectural principles for characterizing the performance of antithetic integral feedback networks,” iScience, vol. 14, pp. 277–291, 2019.

[16] B. Shannon, et al., “In vivo feedback control of an antithetic molecular-titration motif in escherichia coli using microfluidics,” ACS Synthetic Biology, vol. 9, no. 10, pp. 2617–2624, 2020.

[17] D. Salzano, et al., “Ratiometric control of cell phenotypes in monostrain microbial consortia,” Journal of The Royal Society Interface, vol. 19, no. 192, 2022.

[18] S. M. Brancato, et al., “Ratiometric control of two microbial populations via a dual chamber bioreactor,” IEEE Control Systems Letters, 2024.

